# Optogenetic control of medaka behavior with channelrhodopsin

**DOI:** 10.1101/2023.04.05.535638

**Authors:** Takahide Seki, Hideaki Takeuchi, Satoshi Ansai

## Abstract

Optogenetics enables the manipulation of neural activity with high spatiotemporal resolution in genetically defined neurons. The method is widely used in various model animals in the neuroscience and physiology fields. Channelrhodopsins are robust tools for optogenetic manipulation, but they have not yet been used for studies in medaka. In the present study, we used the CRISPR/Cas9-mediated knock-in approach to establish a transgenic medaka strain expressing the *Chloromonas oogama* channelrhodopsin (CoChR) in the *ISL LIM homeobox 1* (*isl1*) locus. We demonstrated that light stimuli elicited specific behavioral responses such as bending or turning locomotion in the embryos and pectoral fin movements in the larvae and adults. The response probabilities and intensities of these movements could be controlled by adjusting the intensity, duration, or wavelength of each light stimulus. Furthermore, we demonstrated that the pectoral fin movements in the adult stage could be elicited using a laser pointer to irradiate only the hindbrain region. Our results indicate that CoChR allows for manipulation of medaka behaviors by activating targeted neurons, which will further our understanding of the detailed neural mechanisms of social behaviors in medaka.

## 1 Introduction

Medaka fish (*Oryzias latipes*) are small teleost fish that have attracted increasing attention in the field of social neuroscience. The sexually mature medaka female has a 24-h reproductive cycle and spawns every morning (Egami, 1954; Takahashi et al., 2013), making it possible to assess social behaviors using female medaka under synchronized reproductive conditions. In addition, medaka exhibit various social behaviors derived from their high visual social cognitive skills, such as individual/familiarity recognition (Fukamachi et al., 2009; Imada et al., 2010; Nakayasu & Watanabe, 2014; Okuyama et al., 2014; Wang & Takeuchi, 2017). Molecular genetics tools, such as genome editing (Ansai & Kinoshita, 2014; Watakabe et al., 2018) and efficient spatiotemporal control of gene expression such as by the Tet-On system (Hosoya et al., 2021), are available for medaka and allow for detailed dissection of molecular and neural functions. By taking advantage of these features, studies of the molecular and neural bases of social behaviors have been performed using medaka fish (Okuyama et al., 2014; Shimmura et al., 2017; Kagawa et al., 2017; Yokoi et al., 2020).

Optogenetics allows for the manipulation of neural activity with high spatiotemporal resolution in genetically defined neurons. Optogenetic actuators, light-gated channels and pumps including channelrhodopsin, are used for the manipulation (Adamantidis et al., 2015; Boyden, 2011; Deisseroth & Hegemann, 2017; Del Bene & Wyart, 2012; Häusser, 2014; Miesenböck, 2009). The application of optogenetic actuators in the medaka brain will greatly contribute to our understanding of how subsets of neurons process visual information in social recognition (Okuyama et al., 2017). Although optogenetic tools are widely and effectively used in zebrafish (Del Bene & Wyart, 2012; Häusser, 2014; Portugues et al., 2013; Varady & Distel, 2020), they have not yet been applied to medaka. In the present study, using CRISPR/Cas9-mediated knock-in via non-homologous end joining, we generated transgenic medaka expressing channelrhodopsin (CoChR) in specific cell populations and demonstrated that the optogenetic tool can be used to control behaviors in medaka.

## 2 Materials and methods

### 2.1 Fish

The hi-medaka strain provided by the National BioResource Project medaka (MT835; https://shigen.nig.ac.jp/medaka/) was used as the parental strain of the transgenic fish. The fish were maintained in an aquarium with recirculating water at 26–28°C illuminated by white fluorescent tubes on a 14/10-h light/dark cycle. The fish were fed *Artemia* sp. nauplii once per day and commercial food pellets (larva: Medaka no Mai Baby, Kyorin; juvenile: Medaka no Mai Next, Kyorin; adult: Otohime B2, Marubeni Nisshin Food). Animal handling and experimental procedures were approved by the animal care and use committee of Tohoku University (permission number: 2022LsA-003).

### 2.2 Construction of donor DNA for knock-in

The channelrhodopsin gene from *Chloromonas oogama* (CoChR) was amplified from pTol1-UAS:CoChR-tdTomato (Addgene Plasmid: 124233) (Antinucci et al., 2020) and fused in-frame with a P2A sequence (Inoue et al., 2016) followed by mScarlet with a point mutation (513C>A) to delete the internal NotI site. This expression cassette was cloned into the KpnI/NotI site of a knock-in donor vector, pUC-BaitD-Xhbb-GFP (Kayo et al. in prep). The resulting construct, pUC-BaitD-Xhbb-CoChR-P2A-mScarlet, was extracted using a standard alkaline lysis method, treated with proteinase K (0.4 µg/µL) and 0.5% sodium dodecyl sulfate at 55°C for 30 min, and then purified using NucleoSpin Gel and PCR Clean-up kit (MACHEREY-NAGEL) with Buffer NTB.

### 2.3 Preparation of Cas9 nuclease and sgRNAs

RNA for the Cas9 nuclease was prepared as previously described (Ansai & Kinoshita, 2014) with some modifications. Capped RNA was synthesized from the pCS2+hSpCas9 vector using an AmpliCap SP6 High Yield Message Maker Kit (Cell Script) and then purified with a Monarch RNA Cleanup Kit (50 µg) (New England BioLabs).

An sgRNA targeting site (5’-AGCATGACGCAGCAGAAACGAGG -3’) in the 5’ upstream region of medaka ISL LIM homeobox 1 (*isl1*; Ensembl gene ID: ENSORLG00000028606) was selected using CCTop (Stemmer et al. 2015). Similarly, sgRNA-slc2a15b_exon2-T1 (5’-CCTGGCAGTCGTCAACTCACCTG -3’) and sgRNA-slc2a15b-exon12-T17 (5’-ATGAACGGATATGGAACTCTTGG -3’) were designed for targeted disruption of *slc2a15b*. A 57-mer oligonucleotide containing a T7 promoter sequence and an 18-mer custom-designed target sequence (5’-TAATACGACTCACTATAGG[N_18_]GTTTTAGAGCTAGAAATAGC -3’) was synthesized for each sgRNA target or the BaitD sequence (Murakami et al. 2017). Each sgRNA template was synthesized by polymerase chain reaction (PCR) with 400 nM of the 57-mer custom-designed oligonucleotide, 4 nM of an 80-mer reverse oligonucleotide for the sgRNA 3’ sequence (5’-GTTTTAGAGCTAGAAATAGCAAGTTAAAATAAGGCTAGTCCGTTATCAACTTGAAAAAGTGGCACCGAGTCGGTGCTTTT -3’), and 400 nM of a 20-mer reverse primer for the 3’ end of the sgRNA sequence (5’-AAAAGCACCGACTCGGTGCC -3’). The PCR products were amplified using PrimeSTAR GXL DNA polymerase (Takara Bio) under the following conditions: 35 cycles of 10 s at 98°C, 15 s at 55°C and 30 s at 68°C, and then purified using NucleoSpin Gel and PCR Clean-up (MACHEREY-NAGEL). The purified product (100 ng) was used as a template for *in vitro* transcription and purification using CUGA7 gRNA Synthesis Kit (Nippon Gene).

### 2.4 Establishment of the transgenic strain

An injection mixture containing 25 ng/μl sgRNAs for the BaitD and *isl1* sequences, 100 ng/μl Cas9 mRNA, and 5 ng/μl donor plasmid was injected into 1-cell stage medaka embryos. The injected fish were backcrossed with wild-type (WT) fish, and then F_1_ fish harboring red fluorescent protein (RFP) fluorescence were identified under an SZX16 fluorescence stereomicroscope (Olympus). The RFP-positive fish were genotyped by PCR amplification with the primers isl1-RV1 (5’-CGAGGAGGGAACGTAAACAA -3’) and Xhbb-RV-KpnI (5’-ACCGGTACCGCCAAAGTTGAGCGTTTATTC -3’). The PCR amplicons were analyzed using an automated electrophoresis system MultiNA MCE-202 (Shimadzu). The F_2_ fish generated by backcrossing the F_1_ fish were used for the behavioral analyses.

### 2.5 Fluorescence observations

To remove leucophore pigmentation for optical imaging, 25 ng/μl each of slc2a15b-sgRNA#1 and slc2a15b-sgRNA#2 was injected with 100 ng/μl of Cas9 mRNA into the fertilized eggs of *Tg(isl1-Xhbb:CoChR-2A-mScarlet)*. After hatching, at 11–13 days post-fertilization (dpf), the injected larvae with little pigment cells were selected under an Olympus SZX16 fluorescence stereomicroscope. The larvae were anesthetized with 0.03% ethyl 3-aminobenzoate methanesulfonate (MS-222) (MilliporeSigma) and then mounted in 2% low-melting point agarose (Agarose-LM, Nacalai Tesque) with 0.03% MS-222.

Imaging was performed using an Axio Zoom.V16 (Zeiss) with a green fluorescence protein (GFP) filter set at 38 HE, RFP filter set at 63 HE, the Apotome 3 system, and a monochrome Axiocam 705 mono camera. The images were acquired, compiled into a single image using the Stitching function, and then processed into maximum projections and adjusted brightness, contrast, and gamma using Zen 3.3 Pro (Zeiss).

### 2.6 Optogenetic stimulation test

The fish were placed differently on dishes for testing at each stage. For the embryonic assay, embryos at 6 dpf were placed into grooves (1.5 mm wide, 1.5 mm deep) on 2% agarose. For the larval assay, larvae at 15–20 dpf were anesthetized using 0.03% MS-222 and then individually embedded on 2% low-melting point agarose on a dish. Agarose gel covering the tail was removed to allow the tail to move freely. For the adult assay, fish (>3 months after hatching) were placed into grooves (10 mm wide, 35 mm long, 7 mm deep) on 2% agarose filled with breeding water. Because the testing fish were not fixed, each behavioral assay was initiated when the fish were in a resting state.

Behavioral responses were monitored under a stereomicroscope (SZX16, Olympus) with a high-speed camera (HAS-U1, DITECT) at 500 fps for the embryos and larvae or 200 fps for the adults under LED illumination (SZX2-ILLTS, Olympus) with a high-pass filter (Lee Filters #106, Primary Red). Optogenetic stimulation light was irradiated from above over the whole body of the medaka using the X-Cite XYLIS LED illumination system (XT720S, Excelitas Technologies) with filtering by a 562-(excitation: FF01-562/40-25, emission: FF02-641/75-25, Opto-Line) or 470-nm (SZX2-FGFP, Olympus) filter. The LED illumination was controlled by a TTL serial connection with Raspberry Pi 3 model B (Raspberry Pi Foundation). To determine the time of onset of each light stimulus, a sheet marked with a highlighter (Textsurfer Gel Orange, STAEDTLER) was placed under the dish. Irradiance of each stimulus was measured using a spectrometer (light analyzer LA-105, NK system). Heterozygous KI fish harboring both the KI and WT alleles of *isl1* and their control siblings without the KI allele were obtained by backcrossing KI heterozygous fish with WT fish. To acclimate the testing fish to the light environment, the fish were placed on the stage only under the transmitted light for at least 30 s prior to the behavioral experiments.

### 2.7 Laser stimulation test

Transgenic (Tg) adult fish were placed into grooves (10 mm wide, 35 mm long, 7 mm deep) on 2% agarose filled with the breeding water. A transparent acrylic plate was placed 120 mm above the agarose plate. The medaka was stimulated with blue light (450 nm) at an intensity of 49.9 μW/mm^2^ using a laser pointer (SPB1, Up’nC) manually over the acrylic plate. The behavioral response of each fish was recorded at 60 fps using a digital camera (SW5069Pro, ShenZhen YANGWANG Technology Co., Ltd) with a macro lens (AI Nikkor 28mm f/2.8S, Nikon) over the acrylic plate. Irradiance of the light stimulus was measured using a spectrometer (light analyzer LA-105).

### 2.8 Behavioral data analysis

The behavioral data was processed using Fiji of ImageJ (Schindelin et al., 2012). For analysis of embryos, a region of interest (ROI) encompassing each embryo was specified manually using the ‘Oval’ tool. To determine the timing of the onset and offset of each light stimulus, the area colored by fluorescent highlighting was specified as another ROI. In each ROI, the absolute difference from the previous frame (Δpixel) was calculated. The presence or absence of the behavioral response of each embryo was determined automatically by the following method. First, from the frame with the highest or second highest Δpixel value in the ROI of the highlighting, the frame with the smaller or larger frame number was then designated as the onset or offset frame for each light stimulus, respectively. In the ROI of each embryo, the frames corresponding to the light onset and offset as well as the frames immediately before and after (onset ± 1 and offset ± 1 frames), were excluded from the following analyses. When the duration of each stimulus was ≤250 ms, the mean (µ_1_) and standard deviation (σ_1_) of the Δpixel values of 100 frames from the end of each video in the embryo ROI was calculated. Each embryo with at least 1 frame with a Δpixel value higher than the threshold (µ_1_ + 9 * σ_1_) after the onset frame was defined as an embryo with a behavioral response. When the duration of each stimulus was 500 ms or 1000 ms, we calculated the mean (µ_2_) and standard deviation (σ_2_) of the Δpixel values using the duration with the smaller variance of the 50 frames either immediately before offset or after the onset of each stimulus. The value µ_1_ + 9 * σ_1_ was used as a threshold during the light irradiation and the other value µ_2_ + 9 * σ_2_ was used as a threshold after the stimulus. For the larval and adult analysis, the ‘Angle’ tool was used to measure the differences in pectoral fin angles before and after the light stimulus (Δangle). The fin angle after the stimulus was measured in the frame with maximum fin opening in each fish. The Δangle values in each fish were calculated as averaged values in both pectoral fins.

For the laser stimulation test, pectoral fin angles were measured at the laser-stimulating frame onset and offset and then the Δangles were calculated as described above. The laser-illuminated area of each stimulus was defined using a maximum intensity projection of the frames with light irradiation (including >100 ms for each stimulus), and then the relative position between the edges of the head and tail was calculated as the center of gravity of each illuminated area.

### 2.9 Statistical analysis

All statistical analyses were performed using R version 4.2.0. The effects of irradiance or duration of the light stimuli and genotype (Tg vs WT) on Δangle in the larval and adult tests were examined using the 2-way aligned rank transform (ART) analysis of variance (ANOVA) in the R package *ARTool* version 0.11.1. For the post hoc test, significant differences between all combinations of 2 factors were evaluated by Tukey’s multiple comparison test implemented in the ‘art.con’ function. The relationship between Δangle and wavelength in the larval test was investigated using the Brunner-Munzel test in the R package *brunnermunzel* version 2.0.

## 3. Results

### 3.1 Generation of transgenic medaka expressing a channelrhodopsin

To test whether channelrhodopsins can evoke behavioral responses by light stimulation in medaka, a channelrhodopsin expression cassette was integrated into the *isl1* locus because channelrhodopsin-2 (ChR2) driven by an *isl1* promoter elicits escape behavior in zebrafish (Douglass et al., 2008). We generated a knock-in strain, *Tg(isl1-Xhbb:CoChR-P2A-mScarlet),* by integrating a donor plasmid containing CoChR upstream of the transcriptional start site of the *isl1* locus with a non-homologous end joining-mediated knock-in technique (Figure 1A) (Watakabe et al. 2018). CoChR is reported to have better performance for both behavioral induction and electrophysiologic responses than other blue light-sensitive channelrhodopsins in zebrafish embryos and larvae (Antinucci et al., 2020). This transgenic construct was designed to express RFP (mScarlet) with the CoChR protein in *isl1*-expressing cells by capturing the endogenous enhancer activity.

**Figure 1.**
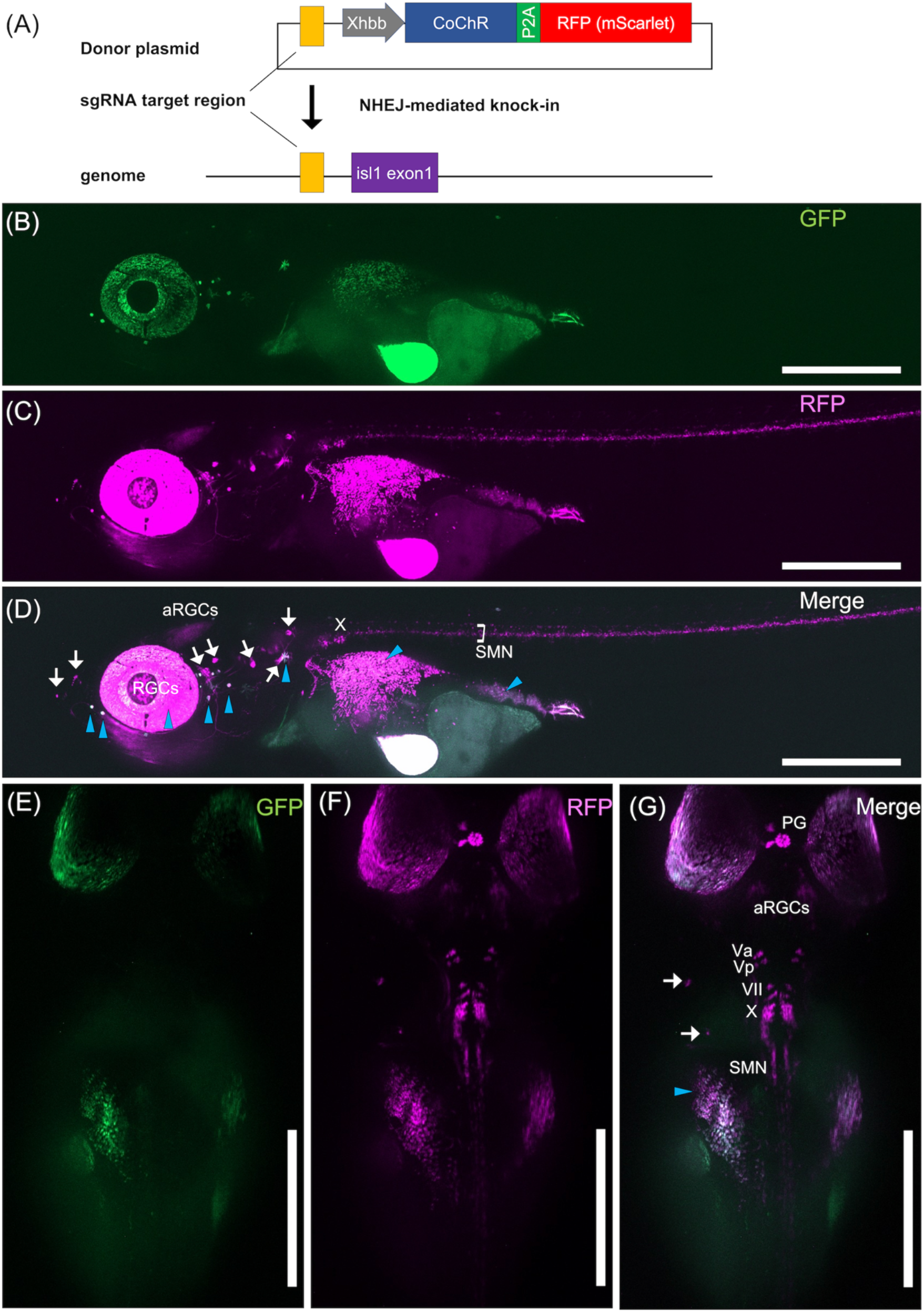
Generation of a transgenic medaka strain *Tg(isl1-Xhbb:CoChR-P2A-mScarlet)*. (A) Schematic illustration of a non-homologous end joining (NHEJ)-mediated knock-in strategy. CoChR, a blue light-sensitive channelrhodopsin from *Chloromonas oogama*, and mScarlet, a red fluorescent protein (RFP), were fused in-frame with a 2A peptide sequence derived from porcine teschovirus (P2A). The expression cassette was driven under the basal promoter derived from the *Xenopus* β-globin gene (Xhbb). Yellow boxes indicate the sgRNA target region. (B-G) Expression patterns of the RFP fluorescence at the larval stage of *Tg(isl1-Xhbb:CoChR-P2A-mScarlet)*. Images with a GFP (B, E) or RFP (C, D) filter set show the autofluorescence or RFP signals, respectively. In the merged images (D, G), white arrows or blue arrowheads indicate RFP signals in ganglions mentioned below or autofluorescence signals, respectively. Both the lateral (B-D) and dorsal (E-G) views are shown. RGCs, retinal ganglion cells; aRGCs, axon of RGCs; V, trigeminal nerve; Va, anterior V; Vp, posterior V; VII, facial nerve; X, vagus nerve; PG, pineal gland; SMN, spinal motor neurons. Scale bars, 500 μm.

In the established transgenic fish, RFP fluorescence was observed in cranial neurons (V; trigeminal nerve, VII; facial nerve, X; vagus nerve) as well as in retinal ganglion cells (RGCs) and their axons (aRGC), spinal motor neurons, the pineal gland, and several sensory ganglions (Figure 1B-G). The *isl1* transgenic zebrafish express GFP in the cranial motor neurons, some of the cranial sensory neurons, and several other groups of cells (Higashijima et al., 2000). The RFP expression patterns of our transgenic medaka mostly matched those of the zebrafish with some differences (such as RGCs or aRGC), suggesting that this transgenic strain successfully expresses the CoChR in *isl1*-expressing neurons. To confirm the activity of CoChR in medaka, we monitored the behavioral responses in several different stages of fish development under light irradiation.

### 3.2 Motor responses induced by optogenetic activation in embryos

We first evaluated whether light activation of the CoChR elicited any behavioral responses in transgenic embryos. Blue light stimulation elicited intense locomotion, such as bending or turning of their bodies, similar to the escape behavior observed in zebrafish embryos (Douglass et al., 2008; Antinucci et al., 2020) (Movie S1). To investigate the effect of blue-light stimulation quantitatively, the optogenetic stimulation test was conducted using the Tg and WT (negative siblings) embryos at 6 dpf (Figure 2A). The embryos were stimulated by the excitation illumination of a fluorescence stereomicroscope and their behavioral reactions were recorded using a high-speed camera (Figure 2A). The effects of the parameters of each light stimulus, such as the duration (5–1000 ms), irradiance (20.7–301.5 μW/mm^2^), and wavelength (470- or 562-nm), on the behavioral responses were investigated. We judged whether each embryo exhibited a motor response by analyzing the difference in the pixel values with the previous frame (Δpixel) (Figure 2B). The probability of the behavioral response of the Tg embryos increased as the light irradiance increased with a continuous duration (1000 ms) and wavelength (470-nm; Figure 2C, blue line).

**Figure 2.**
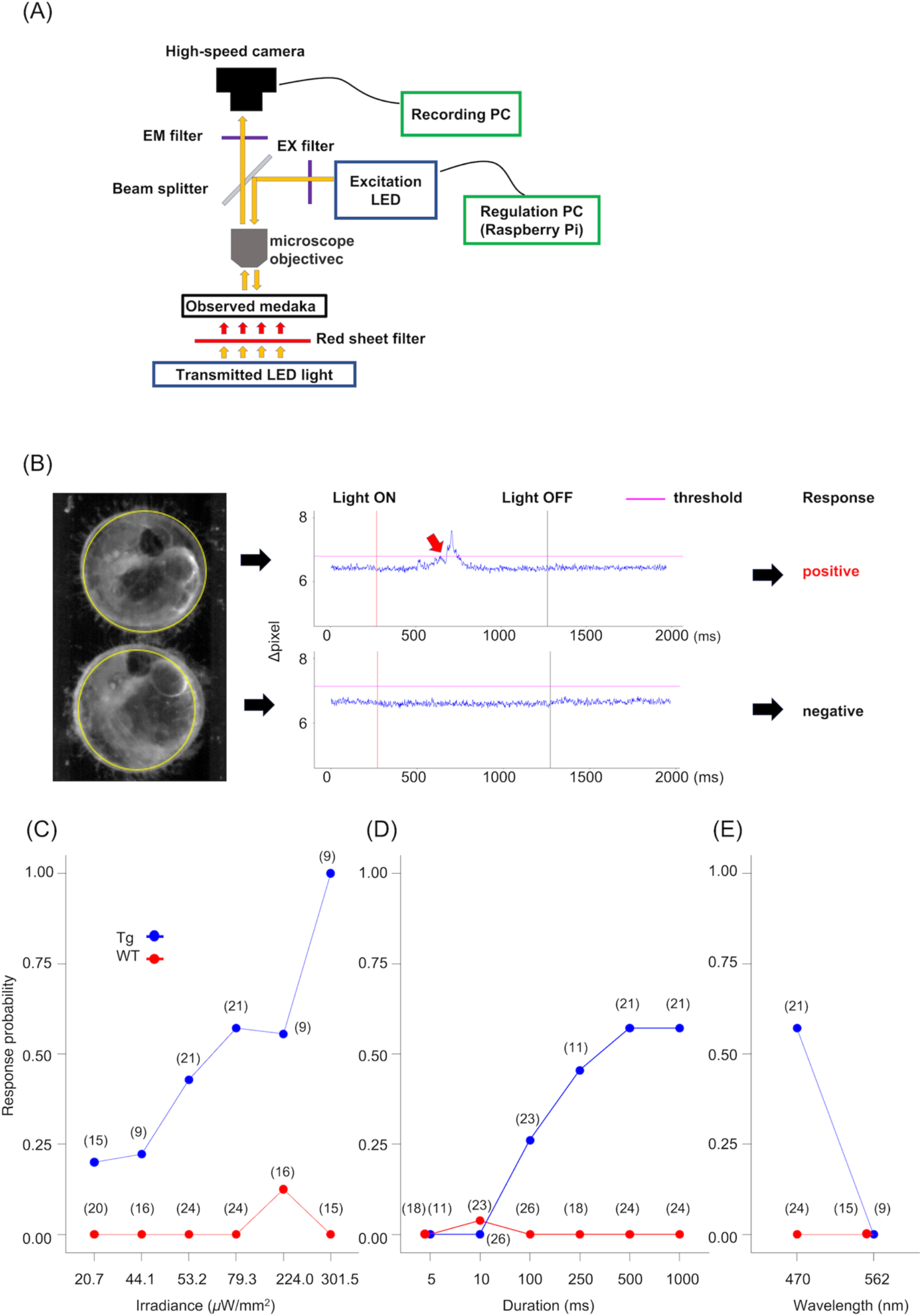
Motor responses to a light stimulus in the embryos of *Tg(isl1-Xhbb:CoChR-P2A-mScarlet)*. (A) Graphical illustration of the experimental setup for the optogenetic stimulation test. (B) Representative traces for behavioral responses triggered by the light stimulus. An embryo shown in the upper panel exhibited a motor response while one in the lower panel exhibited no responses. The behavioral responses were tracked by the difference in pixel values from the previous frame (Δpixel). Red arrow indicates the onset of the response. (C-E) Response probability of the embryos with light stimuli. The blue or red coloration indicates fish harboring the transgenic allele (Tg) or not (WT), respectively. (C) Effects of irradiance on the response probability. Light pulses with a continuous duration (1000 ms) and wavelength (470-nm) were irradiated with different irradiance levels (20.7–301.5 μW/mm^2^). (D) Effect of the duration on the response probability. Light pulses with continuous irradiance (79.3 μW/mm^2^) and wavelength (470-nm) were irradiated with different durations (5–1000 ms). (E) Effects of the wavelengths on the response probability. The 470- or 562-nm light stimulus with 1000-ms duration was irradiated with 79.3 or 245.5 μW/mm^2^, respectively. Sample sizes are shown in parentheses.

Stimulation with a 301.5-μW/mm^2^ light stimulus induced behavioral responses in all of the examined Tg embryos (Figure 2C, blue line). On the other hand, WT embryos rarely exhibited behavioral responses under irradiance at any light intensity (Figure 2C, red line). We also examined the effect of the stimulus duration with a continuous irradiance (301.5 μW/mm^2^) and wavelength (470 nm) and found that the response probability of the Tg embryo increased from 10 ms to 500 ms and reached a maximum at a 500-ms stimulus (Figure 2D, blue line). The WT embryos exhibited no clear responses to the stimuli (Figure 2D, red line). Comparison of the probabilities of the embryos responding to light stimuli with 562- and 470-nm wavelengths revealed that the 562-nm light stimuli induced no behavioral responses in either the Tg or WT embryos and the 470-nm light stimuli induced behavioral responses of the Tg embryos but not the WT (Figure 2E).

### 3.3 Pectoral fin movements induced by optogenetic activation in larval fish

Considering the behavioral response of the Tg embryos, blue-light irradiation was also expected to elicit escape or other swimming behavior in the larvae. Pectoral fin openings, however, were observed as one of the most stable and reproducible responses elicited by blue-light irradiation (Movie S2). To investigate the relationships between the pectoral fin movements and the light stimulus, larval responses to the light stimulus were analyzed using the same setup for optogenetic simulations as in the embryos (Figure 2A). The light stimuli were administered to head-fixed larvae (Movie S3) while changing the irradiance, duration, and wavelength of each light stimulus, similar to the test in embryos. The differences in pectoral fin angles before and after the stimulus (Δangle) were quantified as an index of the behavioral responses (Figure 3A). Both the irradiance of each light stimulus (2-way ART ANOVA, *F* = 4.27, df = 3, *P* < 0.00838) and the genotype (Tg or WT; *F* = 150.27, df = 1, *P* < 0.001) significantly affected the Δangle of the larvae. In the Tg fish, the Δangle increased as the irradiance increased (Tukey’s post hoc test, *P* < 0.00595; Figure 3B, blue). Also, both the duration (2-way ART ANOVA, *F* = 46.94, df = 5, *P* < 0.001) and genotype (*F* = 247.59, df = 1, *P* < 0.001) significantly affected the Δangle values. In the Tg fish, the Δangle significantly increased from the 10-ms stimulus and reached the maximum with a 100-ms stimulus (Tukey’s post hoc test, *P* < 0.05; Figure 3C, blue). The WT larvae did not open their pectoral fins with light stimuli at any irradiance or duration (Tukey’s post hoc test, *P* < 0.05; Figure 3B-C, red). Furthermore, a 562-nm light stimulus with 245.5 μW/mm^2^ irradiance and a 1000-ms duration was irradiated instead of the 470-nm light. Although the Δangle in the Tg fish was significantly increased compared with that in the WT fish (Brunner-Munzel test, *P* = 0.00114; Figure 3D), the Δangle values were smaller than those following the 470-nm stimulus, even with light at a lower irradiance (20.7–79.3 μW/mm^2^; Figure 3B, blue).

**Figure 3.**
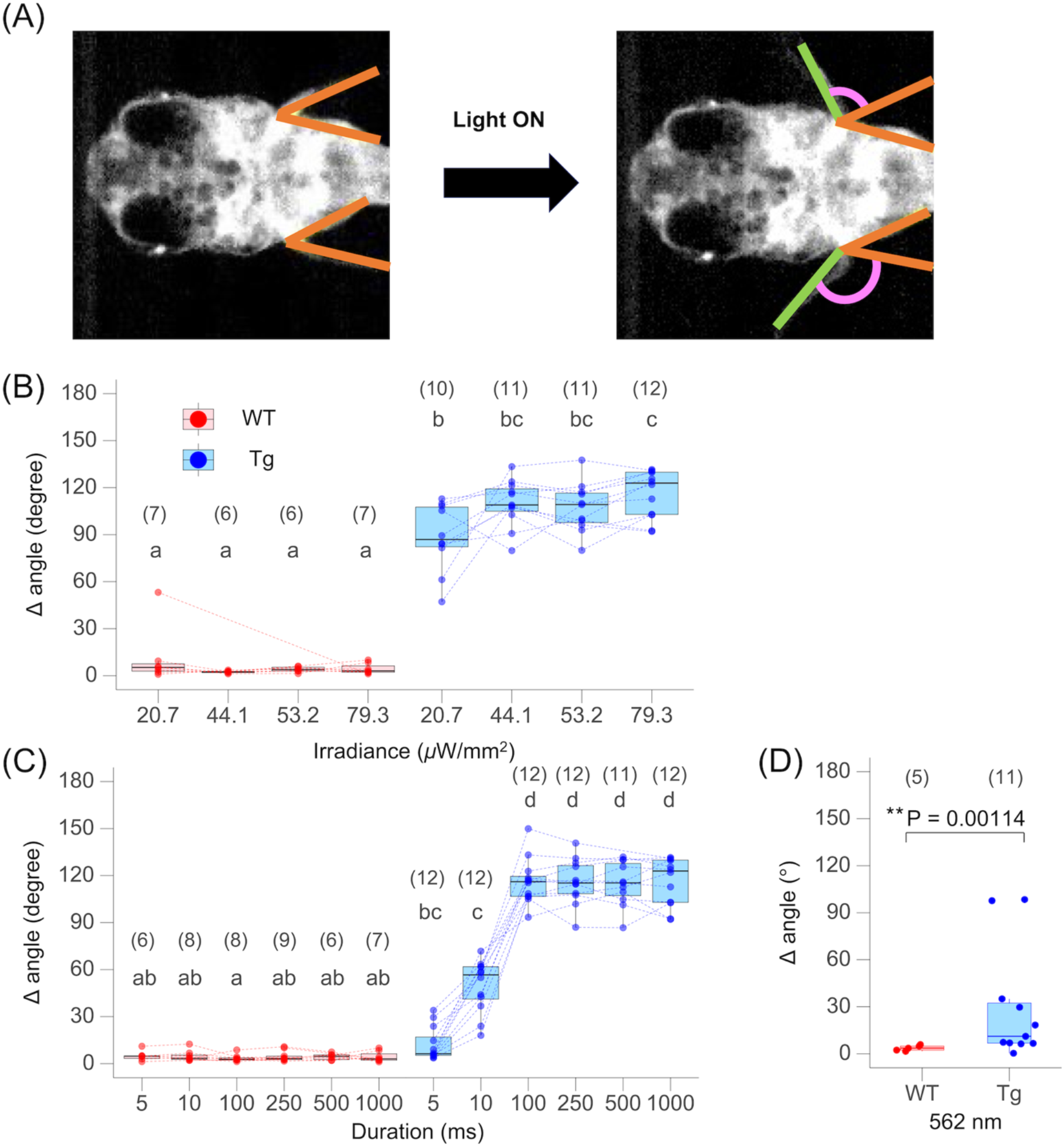
Behavioral responses triggered by a light stimulus in the larval stage of *Tg(isl1-Xhbb:CoChR-P2A-mScarlet)*. (A) A representative illustration for measuring the behavioral responses at the larval stage. Larvae expressing the channelrhodopsin open their pectoral fins after exposure to the light stimulus. Brown and green lines show the angle of the pectoral fins before (left panel) and after (right panel) the light stimulus, respectively. Magenta arcs indicate the difference in the angles (Δangle) before and after the light stimulus. (B-C) Effects of the irradiance (B) or duration (C) of the light stimuli on the Δangle. Boxes show the median, 25^th^, and 75^th^ percentiles for each group. Each dot or line represents an individual. The small letters above each box indicate the results of the ART ANOVA followed by post hoc Tukey’s multiple comparison tests. Different letters indicate significant differences from each other (*P* < 0.05). (B) Light stimuli with a continuous duration (1000 ms) and wavelength (470-nm) were irradiated with different irradiance levels (20.7–79.3 μW/mm^2^). (C) Light stimuli with a continuous irradiance (79.3 μW/mm^2^) and wavelength (470 nm) were irradiated with different durations (5–1000 ms). (D) Effect of the wavelengths on Δangle. The 470- or 562-nm light stimulus with a 1000-ms duration was irradiated with 224.0 or 245.5 μW/mm^2^, respectively. Significant differences were analyzed using the Brunner-Munzel test: \*\**P* < 0.01. Sample sizes are shown in parentheses.

### 3.4 Pectoral fin movements elicited by noninvasive stimuli in adult fish

Finally, we tested whether CoChR works in adult fish. Illumination with blue light also induced pectoral fin openings in adult Tg fish (Figure 4A). Thus, each fish was placed in a small chamber and then illuminated with blue light (Movie S4). We measured the Δangle value of the pectoral fin for each fish and examined the relationships between the Δangle and light stimulus duration (10–1000 ms) or irradiance (9.7–37.0 μW/mm^2^). Similar to the larvae, both the irradiance (2-way ART ANOVA, *F* =17.7, df = 2, *P* < 0.001) and genotype (*F* = 139, df = 1, *P* < 0.001) significantly affected the Δangle value (Figure 4B). In the Tg fish, the Δangle increased as the irradiance increased (Tukey’s post hoc test, *P* < 0.05; Figure 4B, blue). Both the stimulus duration (2-way ART ANOVA, *F* = 21.0, df = 3, *P* < 0.001) and genotype (*F* = 104, df = 1, *P* < 0.001) also significantly affected the Δangle (Figure 4C). In the Tg fish, light stimulation with a duration ≥50 ms significantly increased the Δangle values over that of the WT or Tg fish with a 10-ms stimulus (Tukey’s post hoc test, *P* < 0.05; Figure 4C). On the other hand, the WT fish did not show clear responses under light stimuli of any irradiance or duration (Tukey’s post hoc test, *P* < 0.05; Figure 4B-C, red).

**Figure 4.**
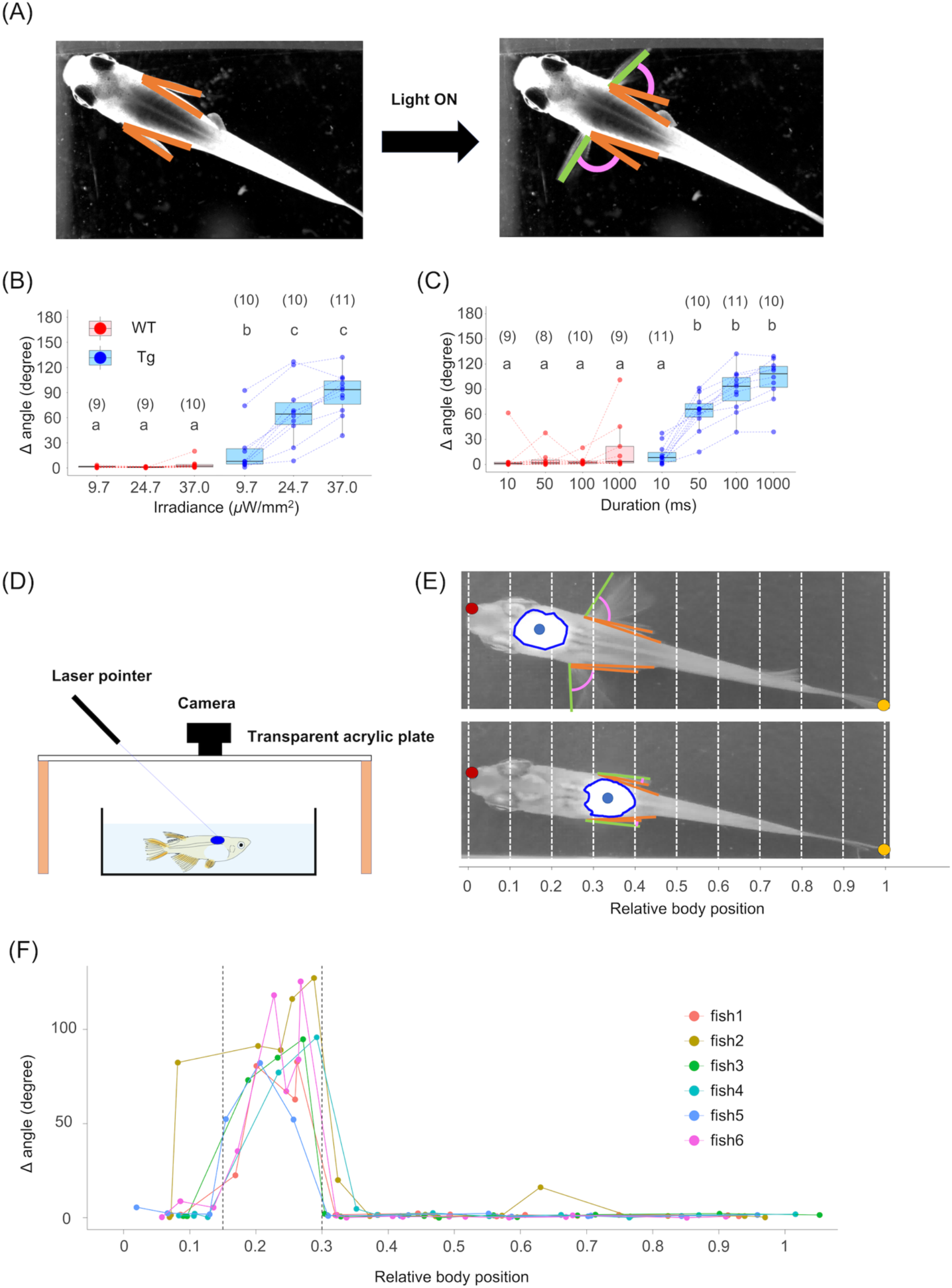
Behavioral responses triggered by light stimulus in adult fish of *Tg(isl1-Xhbb:CoChR-P2A-mScarlet)*. (A) A representative image of the pectoral fin opening observed in adult medaka expressing the channelrhodopsin under a light stimulus. Brown and green lines show the pectoral fin angle before (left panel) and after (right panel) the light stimulus, respectively. Magenta arcs indicate the difference in the angles (Δangle) before and after the light stimulus. (B-C) Effects of the irradiance (B) and duration (C) of the light stimulus on the Δangle. Boxes show the median, 25^th^, and 75^th^ percentiles for each group. Each dot or line represents an individual. The small letters above each box indicate the results of ART ANOVA followed by post hoc Tukey’s multiple comparison tests. Different letters indicate significant differences from each other (P < 0.05). Sample sizes are shown in parentheses. (B) Relationship between irradiance and Δangle. Light stimuli with a continuous duration (100 ms) and wavelength (470-nm) were irradiated at different irradiance levels (9.7–37.0 μW/mm^2^). (C) Relationship between duration and Δangle. Light stimuli with a continuous irradiance (79.3 μW/mm2) and wavelength (470-nm) were irradiated with different durations (10–1000 ms). (D) Experimental setup of the laser stimulation test. Light stimuli with 49.9 μW/mm2 of the irradiance and 450-nm of the wavelength were irradiated to adult fish of the Tg strain using a laser pointer. (E) A representative image generated by maximum intensity projection of the fish images irradiated by the laser pointer to identify the illuminated area. Red, orange, and blue points indicate the edge of the head, the tail edge, and the center of gravity of each illuminated area, respectively. (F) Effects of the relative position of the illuminated area on the Δangle values (n = 6). Each dot or line represents an individual. Dotted lines indicate the relative positions 0.15 and 0.3.

### 3.5 Pectoral fin movements elicited by stimulating only the hindbrain region

The Tg fish were expected to express the CoChR protein in multiple neuronal subsets as observed by RFP expression (Figure 1B-G). To investigate which subsets of the neurons could trigger the pectoral fin movements, adult fish were irradiated locally using a blue-light laser pointer (Figure 4D; Movie S5). The center of gravity of each laser-hitting region was calculated based on the maximum intensity projections of each irradiation, and then the relative body position of each point was determined (Figure 4E). The behavioral responses could be elicited by local irradiation in the 0.15–0.30 region (Figure 4F, between dot lines), in which the hindbrain was included (Figure 4E). Irradiation at positions outside the 0.15–0.30 region did not induce the fin movements (Figure 4F), indicating that the RFP-positive subpopulations outside the 0.15–0.30 region such as pineal gland, aRGCs, and spinal motor neurons, were not involved in the behavioral responses.

## 4. Discussion

In the present study, we established a transgenic medaka strain expressing the channelrhodopsin CoChR and performed a quantitative behavioral analysis to investigate whether blue-light irradiation could induce motor responses. Our results clearly demonstrated that CoChR successfully controlled the neuronal activities of the medaka in vivo. First, the Tg fish exhibited clear behavioral responses to light stimuli whereas the WT fish did not. Second, their responsiveness increased with increased light intensity. Third, the 470-nm light stimulus induced behavioral responses in the Tg fish, while the 562-nm light induced no responses in the embryos and few responses in the larvae. Based on these results, we demonstrated that the ectopic expression of CoChR on medaka neurons is specifically responsible for the behavioral responses induced by blue-light irradiation. The increased responsiveness to the light intensity is consistent with findings from a previous study in CoChR-expressing zebrafish (Antinucci et al., 2020), indicating that CoChR produces a larger photocurrent at higher light intensities (Ganjawala et al., 2019; Shemesh et al., 2017). Moreover, the weak behavioral responses at 562 nm observed in the larvae may be explained by the possibility that 562 nm light, although outside the absorption peak of CoChR, is in the absorption range and thus can generate relatively low photocurrents (Antinucci et al., 2020; Shemesh et al., 2017). To our knowledge, this is the first application of optogenetic tools to manipulate medaka neurons.

The CRISPR/Cas9-mediated knock-in approach used in this study enabled the expression of CoChR in target cells by selecting the target genes. We also demonstrated that CoChR can be activated by direct light irradiation from outside the body without surgery in adult fish. These findings indicate that functional analysis of specific neuronal populations by the optogenetic system established in this study will deepen our understanding of the molecular and neural bases of a wide range of social behaviors observed in adult medaka fish (Okuyama et al., 2017). An optical imaging system for real-time focused tracking of freely swimming adult medaka fish was recently reported (Sueishi et al., 2020). This technique is likely to enable automatic and precise irradiation of the targeted body parts of adult medaka fish and thus become a robust technique for in vivo optogenetic manipulation to minimize the potential off-target effects on their behaviors. The integration of our optogenetic system with such engineering techniques will promote progress in the field of molecular behavioral neuroscience in fish.

## Acknowledgements

We thank the National BioResource Project Medaka (https://shigen.nig.ac.jp/medaka) for providing the hi-medaka strain (ID: MT835). This work was supported by JSPS KAKENHI (Grant Numbers 21H04773 to H.T. and S.A. and 22H05483 to H.T.), Research Grant in the Natural Science of the Mitsubishi Foundation to H.T., Research Grant in the Life Science of the Takeda Science Foundation to H.T, and Advanced Graduate Program for Future Medicine and Health Care, Tohoku University to T.S.

