## Supplemental Movie legends for "Optogenetic control of medaka behavior with channelrhodopsin"

**Supplemental information**

**Legends to Supplemental Movies**

**Movie S1.** Representative movie of the optogenetic stimulation test in embryos with (Tg) or without (WT) the channelrhodopsin transgenic allele *Tg(isl1-Xhbb:CoChR-P2A-mScarlet)*. A 470-nm light stimulus was irradiated to the 6-dpf embryos at 79.3 $\mu W/mm^2$ irradiance with a 500-ms duration. Two embryos at the top left and one embryo at the top right were Tg, and the others were WT. The area marked with fluorescent highlighting in the lower left indicates when the LED is turned on. Images were captured at 500 frames per second (fps) and played back at 0.1× speed (50 fps).

**Movie S2.** Representative movie showing behavioral responses of transgenic medaka strain *Tg(isl1-Xhbb:CoChR-P2A-mScarlet)* larva under blue-light irradiation. Transgenic larva (15-dpf) were irradiated with a 470-nm light stimulus (126.0 $\mu W/mm^2$ irradiance, 100-ms duration). The area marked with fluorescent highlighting in the lower right corner indicates when the LED is turned on. The larva was observed in the breeding water without fixation. Images were captured at 100 fps and played back at 0.5× speed (50 fps)

**Movie S3.** Representative movie of the optogenetic stimulation test in a transgenic larva *Tg(isl1-Xhbb:CoChR-P2A-mScarlet)*. The larva whose head was fixed by agarose mounting was irradiated with a 470-nm light stimulus (79.3 $\mu W/mm^2$ irradiance, 100-ms duration). The area marked with fluorescent highlighting at the bottom indicates when the LED is turned on. Images were captured at 500 fps and played back at 0.1× speed (50 fps).

**Movie S4.** Representative movie of the optogenetic stimulation test in an adult fish of the transgenic strain *Tg(isl1-Xhbb:CoChR-P2A-mScarlet)*. The adult fish was placed in an agarose chamber and irradiated with a 470-nm light stimulus (37.0 $\mu W/mm^2$ irradiance, 100-ms duration). The area marked with fluorescent highlighting at the bottom indicates when the LED is turned on. Images were captured at 200 fps and played back at 0.1× speed (20 fps).

**Movie S5.** Representative movie of the laser stimulation test in an adult fish of the transgenic strain *Tg(isl1-Xhbb:CoChR-P2A-mScarlet)*. The adult fish was locally irradiated using a laser pointer (450-nm wavelength, 49.9 μW/mm$^2$ irradiance, and >100-ms duration). Images were captured at 60 fps and played back at 1× speed (60 fps).
